# Evolution and comparative genomics of tick-associated *Francisella* endosymbionts: insights into metabolic pathways and historical biogeographic patterns

**DOI:** 10.1101/2025.07.01.660216

**Authors:** Juan S. Echeverry-Pérez, Michele Castelli, Sebastián Muñoz-Leal, Santiago Nava, Davide Sassera, Alberto Sánchez-Vialas, A. Sonia Olmeda, Félix Valcárcel, Juan E. Uribe

## Abstract

Ticks (Ixodida) are the second most important vectors of infectious diseases in vertebrates, after mosquitoes. They also maintain mutualistic relationships with bacteria, such as endosymbionts that provide essential B vitamins absent in their blood diet. The most studied endosymbionts belong to the genera *Coxiella*, *Midichloria*, and *Francisella*. *Francisella* includes endosymbionts (FE), pathogens (FP), putative intermediates (FI), and free-living (FL) strains, making them valuable for evolutionary and comparative genomics. In this study, total DNA from six adult female ticks of the genera *Hyalomma* and *Amblyomma* was sequenced to obtain new FE genomes. Additionally, two deep metagenomes from public data were assembled, resulting in a dataset of 22 *Francisella* strains. This dataset was used to reconstruct a phylogenomic framework and compare vitamin biosynthesis and virulence pathways. An MLST-based dense phylogeny was also built to explore biogeographic patterns. The resulting phylogenomic tree shows FE form a monophyletic group derived from FP, possibly due to historical biogeography or recent horizontal transfers. Comparative analyses reveal that FE retain key metabolic pathways while losing nonessential ones, reflecting a selective genome reduction. These results advance our understanding of symbiont evolution in a changing world, revealing molecular adaptations that underpin tick-symbiont relationships and offering genomic insights with potential applications for disease control.

## Introduction

There are currently just over 1000 tick species considered valid, which are distributed in three families of extant species: Argasidae (≈223 species), Ixodidae (≈790 species) and Nuttalliellidae (1 species) (Mans et al. 2019; Guglielmone et al. 2021; Mans et al. 2021; Guglielmone et al. 2023; Muñoz-Leal et al. 2023; Robbins et al. 2025).They are vectors of multiple infectious diseases, including Lyme disease, Crimean-Congo hemorrhagic fever, tularemia, and anaplasmosis(Farlow et al. 2005; Baneth 2014; de la Fuente et al. 2023), posing a significant One Health challenge (Hayman et al. 2023; Machtinger et al. 2024; Pustijanac et al. 2024). Like most animals, ticks harbor a diverse microbiome consisting of pathogenic, commensal, and mutualistic bacteria(Špitalská et al. 2018; Varela-Stokes et al. 2018; Paez-Triana et al. 2023). These microbes can be transmitted horizontally between ticks or vertically to their offspring (Krawczyk et al. 2022; Bakker et al. 2023; Du et al. 2023), with vertical transmission being key for endosymbionts such as *Coxiella* (CE) (Duron, Noël, et al. 2015), *Francisella* (FE) (Baldridge et al. 2009), *Rickettsia* (RE) (Driscoll et al. 2017) and *Midichloria* (Floriano et al. 2023). Other endosymbionts include *Arsenophonus*, *Rickettsiella*, *Wolbachia*, *Lariskella*, *Spiroplasma,* and *Cardinium* (Ahantarig et al. 2013; Duron et al. 2017; Duron 2024). These microorganisms play essential roles in tick growth, molting, immunomodulation, reproduction, and metabolism, colonizing key organs such as the ovaries, Malpighian tubules, salivary glands, and mitochondria (Sassera et al. 2006; Kolo and Raghavan 2023; Zhong et al. 2024).

The hematophagous diet of ticks is rich in proteins, iron, and salts, but deficient in carbohydrates, lipids, and vitamins (Duarte et al. 1999; Lehane 2005; Sterkel et al. 2017). This dietary imbalance drives their reliance on genomic adaptations (Mans et al. 2002; Sojka et al. 2013) and mutualistic interactions with endosymbiotic bacteria, which supply missing metabolic pathways, particularly those involved in B vitamin biosynthesis (Duron and Gottlieb 2020; Buysse and Duron 2021; Adegoke et al. 2022). FE act as obligate symbionts in multiple tick species, thanks to intact biosynthetic pathways for B2 (riboflavin), B7 (biotin), and B9 (folate) that have been identified in their genomes (Gerhart et al. 2016; Duron et al. 2018; Gerhart et al. 2018; Buysse et al. 2021). The same applies to CE (Gottlieb et al. 2015; Smith et al. 2015; Guizzo et al. 2017), whereas RE (Hunter et al. 2015; Driscoll et al. 2017) and *Midichloria* (Olivieri et al. 2019) possess incomplete pathways.

Compared to pathogenic and free-living species, the genomes of endosymbionts have undergone extensive chromosomal rearrangements and genome reduction, due to a combination of processes driven by selection, genetic drift, and mutation (McCutcheon and Moran 2011; Keeling and McCutcheon 2017; McCutcheon et al. 2024). Throughout their evolution, non-essential genes become pseudogenes and are eventually lost, affecting functions such as pathogenicity, motility, and regulation (Moran et al. 2008; Wernegreen 2017). Although a fully circularized FE genome has not yet been obtained, phylogenomic studies indicate that FE originated from the pathogenic *Francisella* clade. This is consistent with their smaller genome size, the loss of virulence genes, and the retention of essential metabolic pathways (Gerhart et al. 2016; Duron et al. 2018; Gerhart et al. 2018; Buysse et al. 2021). On the other hand, CE exhibit a wide range of genome sizes (0.65-1-73 Mb) (Nardi et al. 2021), including even more pronouncedly reduced and potentially less efficient genomes (Smith et al. 2015), suggesting that some species could be replaced by FE (Duron and Gottlieb 2020; Bontemps et al. 2024) in line with the ‘symbiosis rabbit hole’ theory and Muller’s ratchet (Pettersson and Berg 2007; Bennett and Moran 2015).

The present study examines the evolution and biogeography of FE, as well as their relationship with ticks, through a comparative genomic approach. To achieve this, total DNA of six ticks (*Amblyomma* and *Hyalomma* spp.) was sequenced. Additionally, two publicly available deep metagenomes belonging to *Hyalomma marginatum* ticks were included. From these eight samples, FE genomes were assembled, with five of them successfully circularized. The analysis also incorporated genomic sequences from previously published studies, five FE, two FI, six FP, and one FL strain, for a total of 22 *Francisella* genomes analyzed. These strains represent a broad geographic distribution, covering regions of America, the Circum-Mediterranean, and Asia. Our main finding suggests a biogeographic pattern and potential host-switching events among *Francisella* strains, while also supporting the hypothesis that *Francisella* supplements essential nutrients to its tick hosts and exhibits loss of pathogenicity-related genes, highlighting its evolutionary transition towards a symbiotic lifestyle.

## Materials and methods

### Sample Collection, Identification, and DNA Extraction

Six adult female ticks were collected (see information about the collection in Table S1), fixed in absolute ethanol, and stored at -20°C. Morphological identification was performed following taxonomic guides for *Amblyomma* (Nava et al. 2017) and *Hyalomma* (Apanaskevich and Horak 2008; Estrada-Peña et al. 2017). DNA extraction was carried out using the DNeasy Blood & Tissue Kit (Qiagen), following the manufacturer’s protocol, with an extended proteinase K lysis step for 20 hours at 56°C under continuous agitation. Molecular confirmation of the identification was performed using the COI gene following a previously published protocol (Uribe et al. 2024).

### Genome Sequencing and Assembly

For each sample, a Plant and Animal Whole Genome Sequencing (PAWGS) library was prepared and sequenced on an Illumina NovaSeq 6000 platform (150 bp paired-end; the output size of each sample is included in Table S1) by Novogene. Quality was assessed using FastQC (Andrews 2017), and low-quality reads and adapters were removed with Trimmomatic v0.39 (Bolger et al. 2014). *de novo* assemblies were generated using SPAdes 4.0.0 (Prjibelski et al. 2020) with default parameters.

In order to select the sequences of the FE with respect to tick host and its microbiome, a fine-tuned multi-step procedure was developed and applied to the six newly sequenced samples and to two additional sets of *H. marginatum* tick metagenomic reads from NCBI (Run accessions ERR7255744 and ERR9924217, described in detail in the Supplementary material). Briefly, 12 *Francisella* reference genomes (three FE, eight FP, and one FL; Table S2) were downloaded from NCBI and used as a reference database for a blastn search of the contigs of the preliminary assembly of each sample, and contigs with positive hits were thus labeled as FE sequences. The procedure was iteratively repeated by enriching the reference database with the FE contigs progressively identified from all samples, and, finally, for increasing sensitivity, by applying a tblastn search, then verifying the hits by blast on NCBI databases. Moreover, extracted FE reads used for mapping, *de novo* assemblies improvement (joining them as the case of the circularized), and coverage assessment were performed in Geneious® Prime (Kearse et al. 2012). Finally, the assembly quality of the eight newly obtained FE was assessed along with the other *Francisella* (five FE, two FI, six FP, and one FL) using BUSCO (Manni et al. 2021). Additionally, the mitochondrial genome of the samples sequenced here were assembled, filtered, and annotated following a standard methodology (Abalde et al. 2019).

### Gene Annotation and Metabolic Pathway Analysis

Genome annotation was performed with PGAP (Tatusova et al. 2016; Haft et al. 2018) using default parameters, specifying *Francisella* as the target genus. Intact genes and pseudogenes were identified using the ‘Annotate’ command in Pseudofinder (Syberg-Olsen et al. 2022). Metabolic pathways were inferred with BlastKOALA and GhostKOALA (KEGG) (Kanehisa et al. 2016), and additional blastp searches were conducted based on curated gene lists related to vitamin biosynthesis obtained from reference Biocyc Pathways (Karp et al. 2019) and pathogenicity (Duron et al. 2018). Clusters of Orthologous Groups (COGs) were assigned using eggNOG-mapper, with *Gammaproteobacteria* set as the reference taxonomic group (Cantalapiedra et al. 2021).

### Phylogenetic Analysis

A phylogenomic tree was constructed using the 22 *Francisella* genomes. Predicted amino acid sequences, excluding pseudogenes, were used to identify single copy orthologroups with OrthoFinder v2.5.4 (Emms and Kelly 2019). The matrix was prepared following the pipeline described in (Uribe et al. 2022), consisting of: *(i)* Sequence masking with PREQUAL (Whelan et al. 2018) using default parameters; *(ii)* Multiple sequence alignment with MAFFT v7 (Katoh and Standley 2013) using the default algorithm FFT-NS-2; *(iii)* Filtering of phylogenetically informative regions with BMGE v1.12 (Criscuolo and Gribaldo 2010) under default parameters; *(iv)* Concatenation of the aligned genes using Geneious^®^. A maximum likelihood (ML) tree was inferred with IQTREE v2 (Minh et al. 2020), and branch robustness was evaluated using 1000 ultrafast bootstrap (UFBoot) replicates. The best-fit ML model was determined with ModelFinder (Kalyaanamoorthy et al. 2017), based on Bayesian Information Criterion (BIC) (Schwarz 1978). The tree was rooted using the FL strain according to previous studies (Duron et al. 2018).

Additionally, a phylogenetic tree was constructed based on Multi-Locus Sequence Typing (MLST) genes of FE (*16S rRNA, rpoB, groL, ftsZ,* and *gyrB*) (Binetruy et al. 2020). This analysis included the 22 *Francisella* genomes in this study, along with additional gene sequences retrieved from NCBI Nucleotide (Table S3).

A mitogenomic phylogeny of ticks was also generated, incorporating host species associated with *Francisella* from this study, or closely related congeneric species from the same ecoregion for those samples for which corresponding host mitogenome was not available since they were not sequenced in this study (Table S4). Tick mitogenomes were annotated using MITOS (Bernt et al. 2013), with gene boundaries manually curated. Both the MLST and mitogenomic trees (outgroup: *Nuttalliella namaqua*) were inferred using nucleotide sequences, the best-fit evolutionary model for concatenated matrices (Kalyaanamoorthy et al. 2017) and ML methods (Minh et al. 2020), as described above.

## Results

### General Genome Comparisons

Six new tick samples were sequenced, and seven previously available sets of FE were analyzed. From these data, we *de novo* assembled eight FE genomes and successfully circularized five of them (Figure 1). All FE genomes originate from Neotropic, Nearctic, Palearctic (mainly from the Circum-Mediterranean subregion), Afrotropical, and Indomalayan (Figure 1 and 2). The FE genomes exhibit consistent features such as genome sizes ranging from 1.51–1.67 Mb, GC content between 31.2%–31.9%, BUSCO completeness scores of 80–86.4%, and 875–1,062 coding genes (Table S2). Accordingly, we treated the non-circularized assemblies as complete, since their metrics are comparable to those of the circularized genomes. Compared to other *Francisella* strains, both FE and FI display reduced genomic complexity. FE and FI share similar genome characteristics, namely more compact genomes (∼1.5 Mb), slightly lower GC content (∼31.3%) and BUSCO completeness (∼80–87%), as well as a moderate number of coding genes (∼875–1,050). On the other hand, FP and FL strains exhibit larger and more complex genomes, with sizes of 1.82–2.03 Mb, GC content of 32.2–32.9%, BUSCO completeness of 88.6–91%, and 1,386–1,818 coding genes. In contrast, pseudogene content is highest in FE (653–836), intermediate in FI (451–455), and lowest in FP and FL strains (131–422, except in *F. tularensis* subsp. *holarctica* OSU18, which reaches 529) (Table S2; Figures S1 and S2). A particularly notable case is FE-*Amblyomma parvum* (AM04), which harbors a circular contig of approximately 7.7 Kb containing pathogenicity genes (see below) commonly found on plasmids among FP strains.

**Figure 1.**
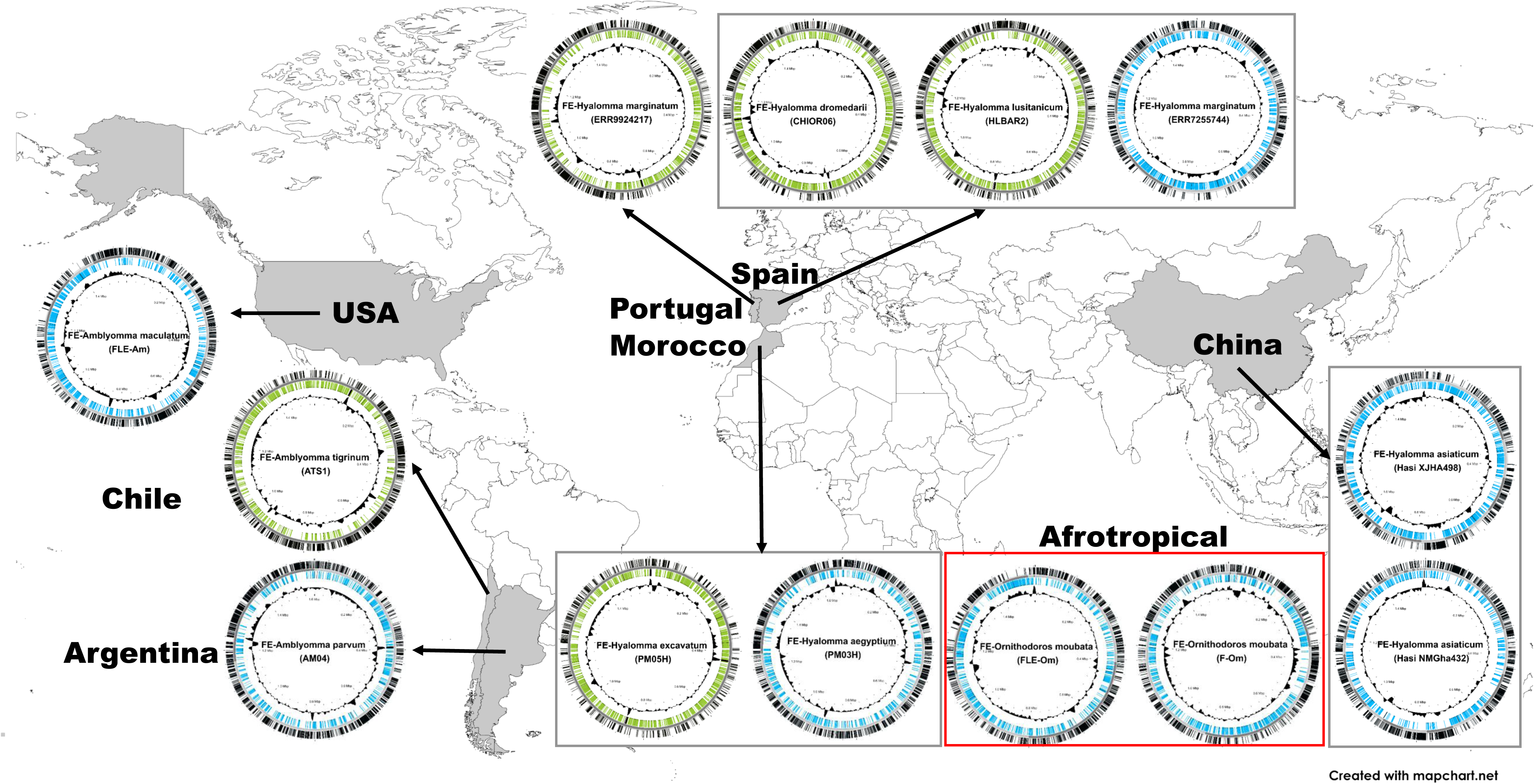
Geographic distribution of *Francisella* endosymbionts (FE) analyzed in this study. Localities of FE samples are mapped, with circularized genomes shown in green and contig-level genomes in blue. The *Hyalomma* group includes eight specimens from six species, five from the Circum-Mediterranean region and one from China. *Amblyomma* is represented by three species from the Americas, while the *Ornithodoros* group from the Afrotropical region includes two specimens of the same species (red inset).

**Figure 2.**
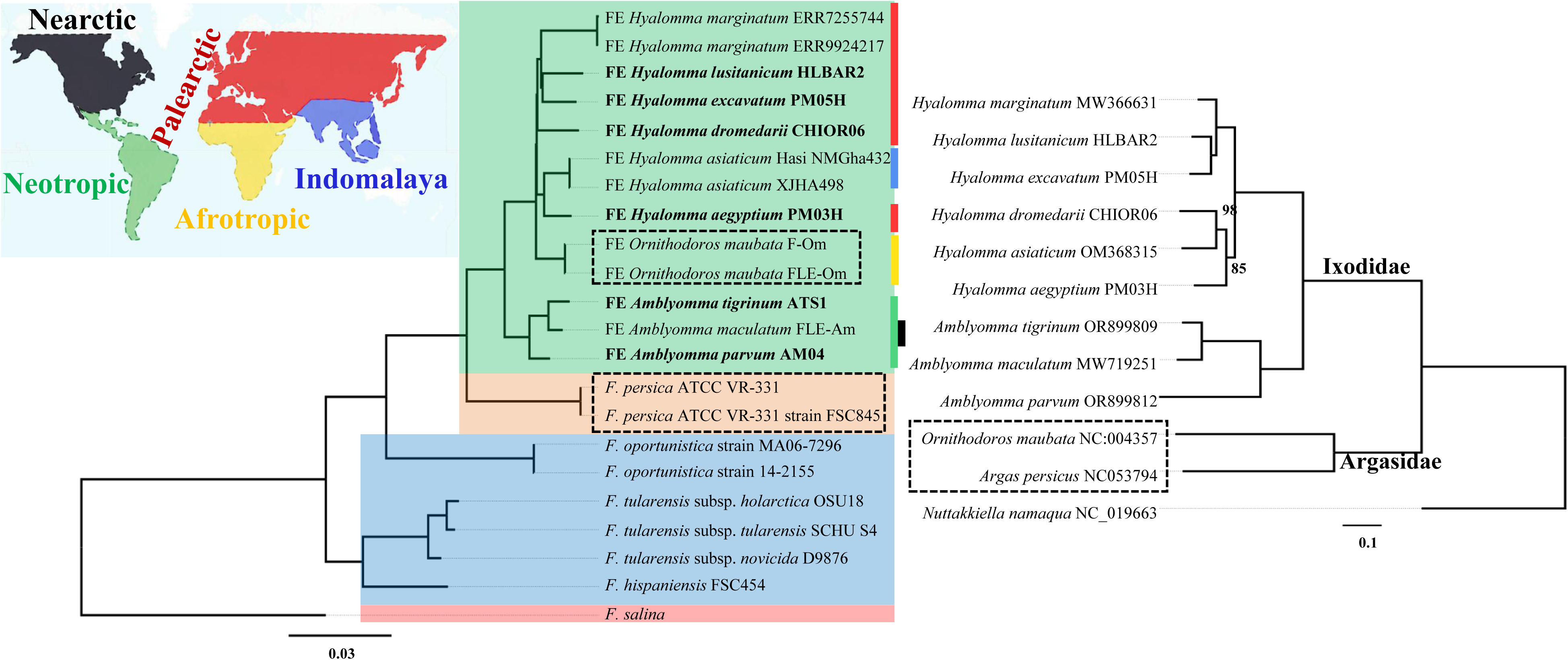
Phylogeny of *Francisella* and its tick hosts. On the left, a maximum-likelihood (ML) phylogenomic tree of *Francisella* based on 429 predicted protein-coding genes. The geographic distribution of each *Francisella* endosymbiont (FE) is indicated with colored side bars corresponding to the map. Background colors denote lifestyle categories: green for FE, peach for FI, blue for FP, and red for the FL strain. On the right, a ML mitogenomic tree of tick hosts, constructed from protein-coding and ribosomal RNA genes at the nucleotide level. *Francisella* strains and their Argasidae tick hosts are highlighted with dotted boxes in both trees. Only nodes with bootstrap support values below 100% are shown. The scale bar represents the expected number of substitutions per site.

### Phylogeny

A maximum-likelihood (ML) phylogenomic tree was constructed using 429 predicted protein sequences from the 22 *Francisella* members analyzed in this study. The topology of the FE tree was compared with the phylogeny of their tick hosts to assess potential coevolutionary processes (Figure 2). The *Francisella* phylogenomic tree exhibits maximum support (Figure 2, left). The topology reveals that pathogenic strains represent the earliest divergences. Within a derived clade, FE are recovered as a monophyletic group, with *F. persica* as its sister clade (Figure 2, left). The FE-*Amblyomma* from America were found as a sister lineage to FE from the rest of the world, including the Circum-Mediterranean, Afrotropical, and Oriental regions (Figure 2A). Within this global distribution, the FE-*Ornithodoros* (Afrotropical) strains appear as a sister clade to the FE- *Hyalomma* strains (Circum-Mediterranean and Oriental). Within the FE-*Hyalomma* lineage, FE-*H. aegyptium* clusters with FE-*H. asiaticum*, forming a sister clade to a group composed of FE-*H. dromedarii*, FE-*H. marginatum*, FE-*H. excavatum*, and FE-*H. lusitanicum*. These phylogenetic relationships within the genus *Francisella* do not fully align with those of their hosts at the host family level, though they do at the host genus level. Indeed, the FE strains of the Argasidae tick *O. moubata* are more closely related to FE of Ixodidae (genera *Hyalomma* and *Amblyomma*) than to *F. persica*, hosted by the Argasidae tick *Argas persicus* (Figure 2, left tree), despite the clear phylogenetic separation of Ixodidae and Argasidae (Figure 2, right tree). Additionally, the FE of hard ticks are also non-monophyletic, since FE of *Hyalomma* spp. are more closely related to the FE of *O. moubata* than to those of *Amblyomma* spp. (Figure 2, left tree).

The MLST tree of *Francisella* (Figure 3; complete tree in Figure S3) reflects a similar pattern of the phylogenomic tree, positioning FE as a derived clade from FP congeners, with FI as a sister group. Within the FE group, the earliest divergence involves the FE*-Amblyomma* from the Neotropical Americas, with the first clade (I) grouping FE hosted by *A. goeldii* and *A. humerale.* This is followed by a second divergence (clade II) of a FE hosted by the Neotropical *A. varium*. The remaining FE strains are splitted into two main sister clades (III and IV). Clade III includes FE hosted both by exclusively Neotropical *Amblyomma* ticks, such as *A. tigrinum*, *A. parvum*, *A. oblongoguttatum*, *A. pacae*, *A. geayi,* and *A. latepunctatum*, as well as FE strains hosted by more widely distributed (Neotropic and Nearctic) *Amblyomma*, such as *A. maculatum*, *A. ovale*, *A. rotundatum*, *A. dissimile,* and *A. longirostre.* Group IV includes FE associated with members of three tick genera, grouped in multiple subclades. Two FE hosted by *Amblyomma sculptum* (Neotropical) and *A. paulopunctatum* (Afrotropical) are sister group of FE associated with *Ornithodoros moubata* (Afrotropical). This mentioned lineage is sister clade of the FE strains hosted by Paelarctic (*Hyalomma marginatum*, *H. lusitanicum*, *H. excavatum*, *H. aegyptium*, *H. dromedarii*) and Indomalayan (*H. asiaticum*) *Hyalomma* ticks. Noteworthily, the MLST tree recovered the same topology of the phylogenomic tree for what concerns the FE strains hosted by *Hyalomma*.

**Figure 3.**
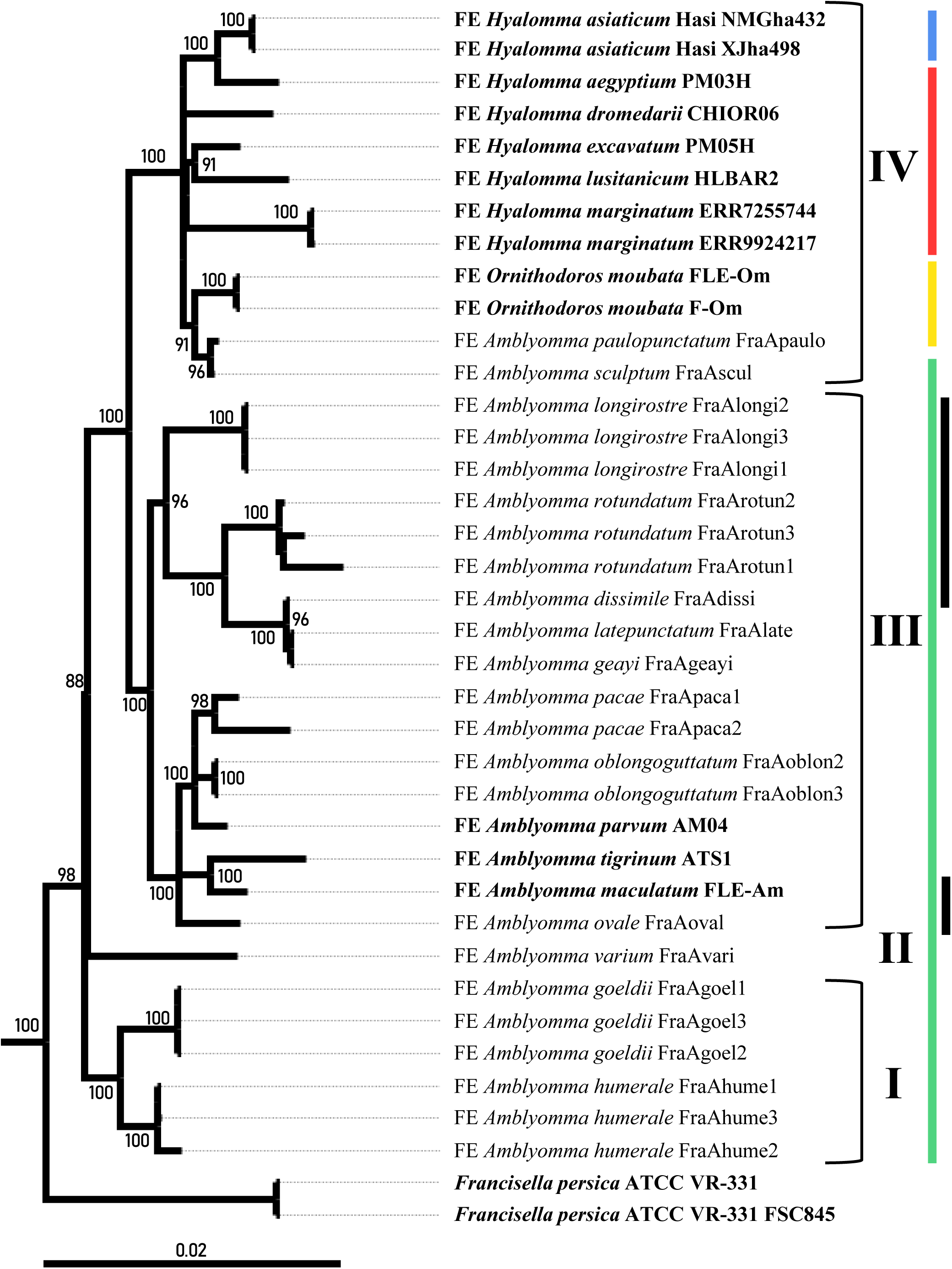
MLST phylogenetic tree of *Francisella* and geographical distribution of tick hosts. The tree reveals four major clades, labeled from bottom to top as Clades I, II, III, and IV. The scale bar indicates the number of expected substitutions per site. Only nodes with bootstrap support values greater than 80% are shown. Colored side bars correspond to the geographical distribution and lifestyle traits shown in Figure 2. *Francisella* strains whose genomes were analyzed in this study are highlighted in bold. Full tree is shown in Figure S3.

### Comparison of Clusters of Orthologous Groups (COG)

The analysis of *Francisella* identified 1,796 unique COGs distributed across 18 COG categories. The number of COGs per genome ranged from 665 in FE-*H. dromedarii* (CHIOR06) to 1,301 in *F. salina* (FL) (Figure 4). Among the endosymbionts, FE-*Amblyomma parvum* (AM04) had the highest number of COGs, with 784, while *F. persica* (FI) strains exhibited 803 and 809 COGs. The pathogenic *Francisella* (FP) ranged from 1022 COGs in *F. tularensis* subsp. *holarctica* OSU18 to 1191 COGs in *F. tularensis* subsp. *novicida* D9876. Across the whole dataset, the most abundant categories in descending order were S (Function unknown). E (Amino acid transport and metabolism). J (Translation, ribosomal structure, and biogenesis), M (Cell wall/membrane/envelope biogenesis), and C (Energy production and conversion). When considering four main groups of *Francisella* members (FE, FI, FP and FL), we found that 792 COG are shared across all groups, while 230 are unique to FP strains, 208 to FL, 45 to FE, and only 1 to FI (Figure 5; more detailed in Figure S5). Given the overall high similarity of the COG content among the FE strains, the following analysis will focus specifically on features previously associated with the host interaction.

**Figure 4.**
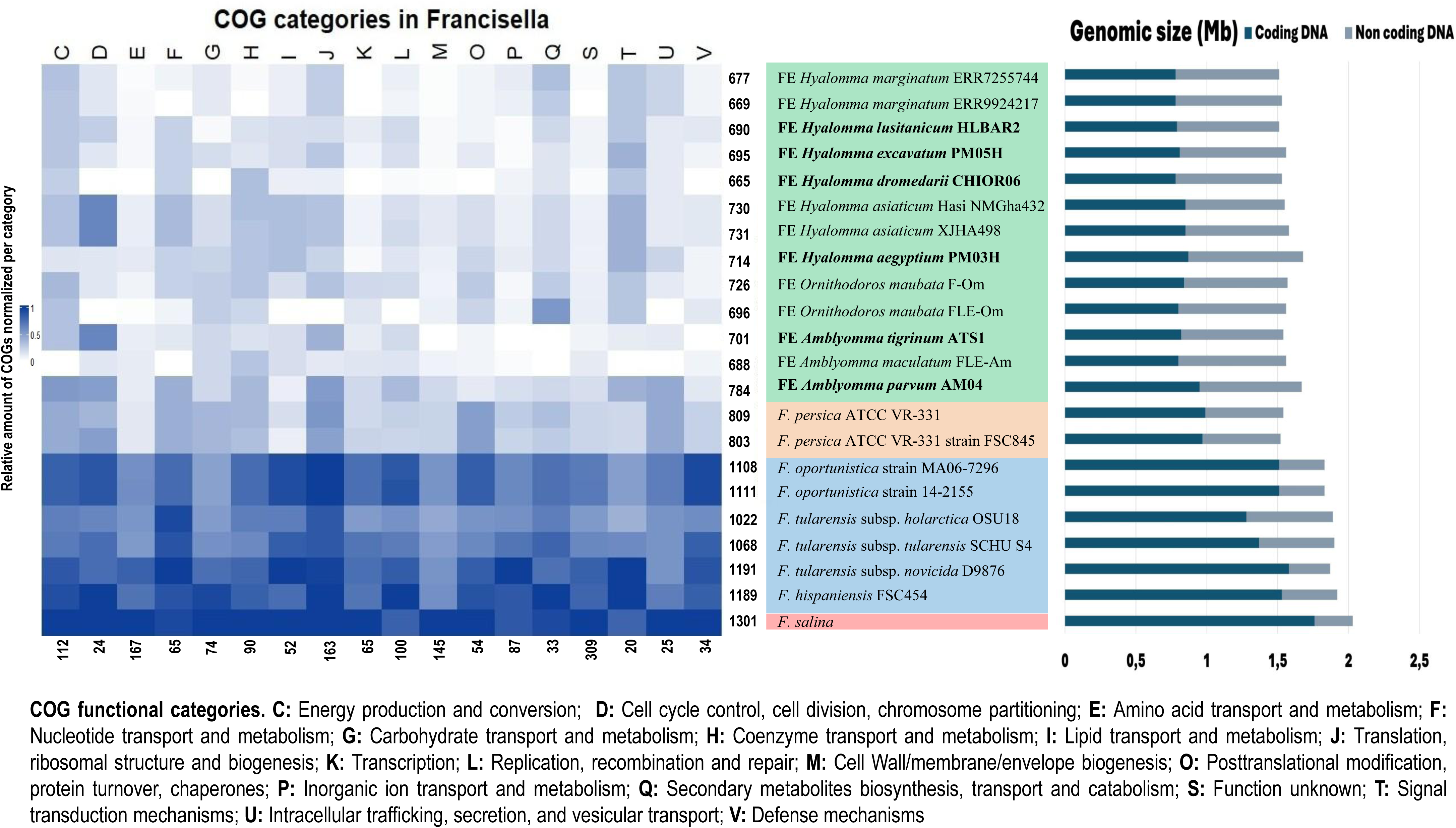
Distribution of Clusters of Orthologous Groups (COG) by category in *Francisella*. At the left of the heatmap, each row represents the total number of COGs per genome, while each column shows the total number of COGs per functional category. In the middle, the organisms are arranged according to the phylogenomic tree from Figure 2. At right, the bar plot represents genome size (Mb), coding and non-coding length of each strain.

**Figure 5.**
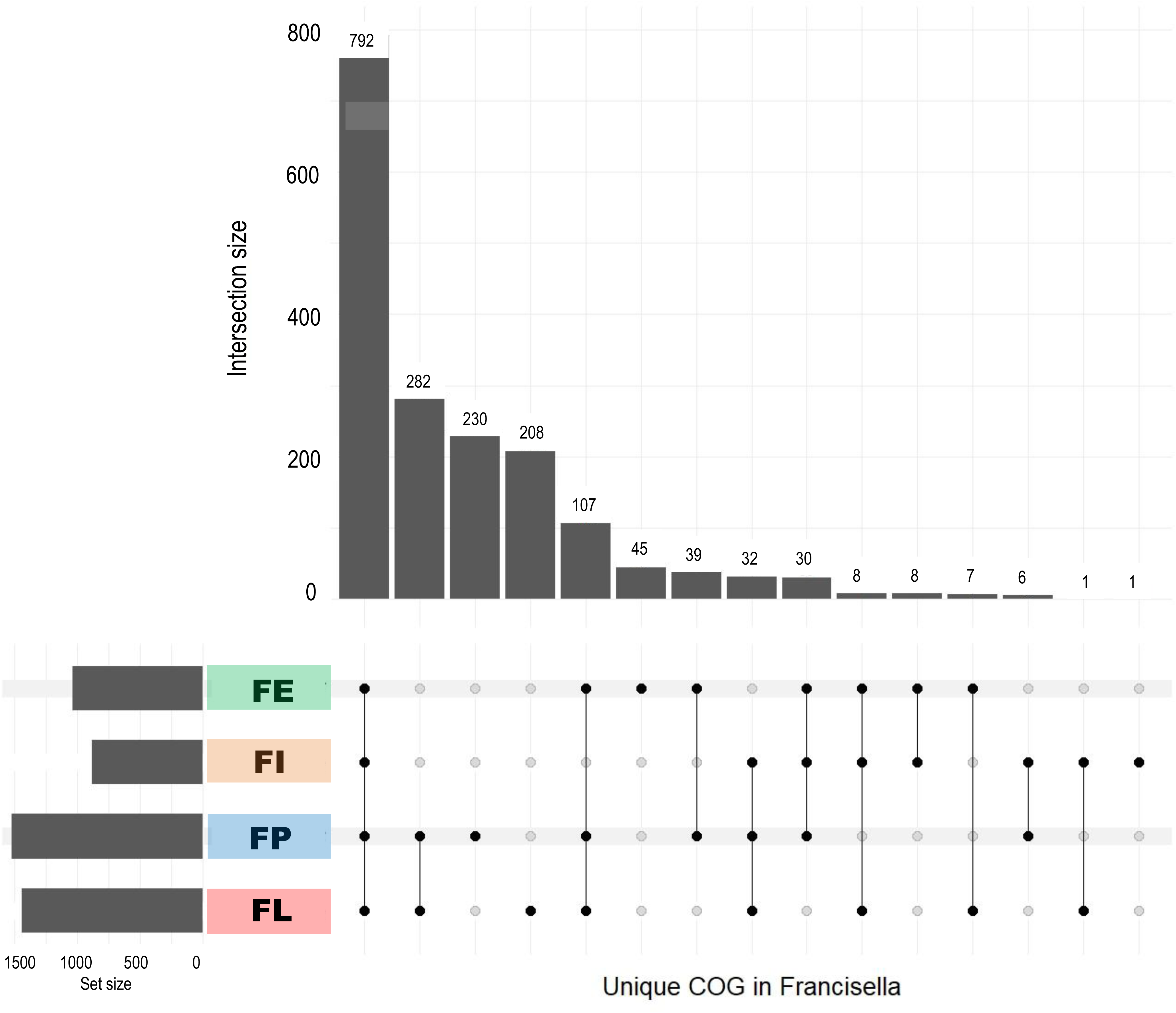
UpSet plot of COG intersections of COGs presence/absence in *Francisella*. The bottom axis displays the *Francisella* groups: FE, FI, FP, and FL. Vertical bars indicate the number of unique COGs shared exclusively among the connected groups.

### Vitamin Biosynthesis Pathways in Francisella

The vitamin biosynthesis pathways in *Francisella* exhibit varying levels of conservation and erosion (Figure 6). Several pathways, such as riboflavin (B2), heme, lipoic acid, and ubiquinone, are broadly conserved across the genus. The synthesis of biotin, recognized as involved in FE strains interactions with ticks (Napier et al. 2012; Feng et al. 2014) is also widely present, although some genes are pseudogenized in a subclade of FE strains from *Hyalomma* spp. (Figure 6). The final steps in the biosynthesis of pantothenate (B5) and Coenzyme A are conserved in all *Francisella* species, with two genes pseudogenized in the FE strains of *O. moubata* (FLE-Om). In contrast, the preliminary steps of this pathway are largely retained in free-living and pathogenic *Francisella*, but are reduced in FE strains, and absent in FI. The main genes involved in pyridoxine (B6) synthesis are present in all *Francisella*, including FE, with the only exception of the two *F. opportunistica (FP)*. Conversely, NAD (B3) biosynthesis is substantially conserved in FP, FL, and FI strains but is either absent or pseudogenized in FE strains. The thiamine (B1) synthesis does not follow a clear pattern within the genus, being found only in FL and *F. hispaniensis*. Finally, the folate pathway also lacks a regular pattern of erosion, displaying random genes absent either in FE, FI, FP and FL strains (Figure 6).

**Figure 6.**
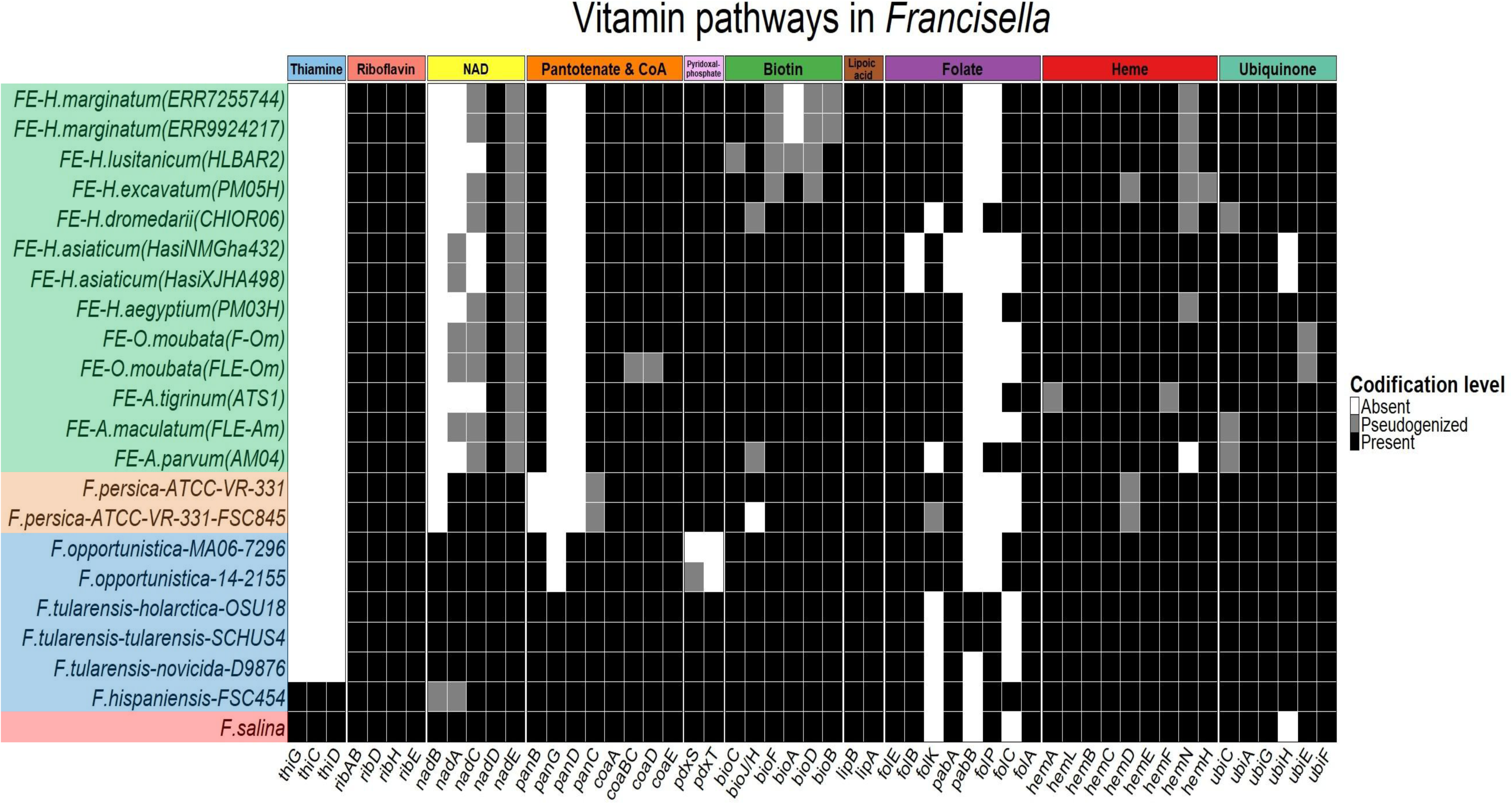
Evolution of vitamin biosynthesis pathways in the *Francisella* genus. The heatmap depicts the conservation of vitamin B and cofactor biosynthetic genes, with a color gradient indicating gene status: dark tones represent intact coding sequences, lighter tones denote pseudogenes, and white indicates gene absence. Each column corresponds to a gene involved in these pathways. The organisms are arranged according to the phylogenomic tree from Figure 2.

### Pathogenicity genes in Francisella

Pathogenicity-related pathways in *Francisella* show extensive variation across the genus, being relatively less represented among FE strains (Figure 7). Putative porins and inner membrane proteins are relatively conserved, but several genes are either missing or pseudogenized in FE strains. Likewise, pathways associated with lipopolysaccharide (LPS), capsule and membrane synthesis, outer membrane proteins, type IV pili, secretion systems, and the *Francisella* pathogenicity island (FPI) are highly eroded in most FE strains, with the majority of genes either pseudogenized or entirely absent. In contrast, these pathways remain at least partially intact in FP and FL strains. Notably, more genes are preserved in FI and in FE-*Amblyomma parvum* (AM04) than among the other FE, such as those related to type IV pili and secretion systems (the latter particularly in FI).

**Figure 7.**
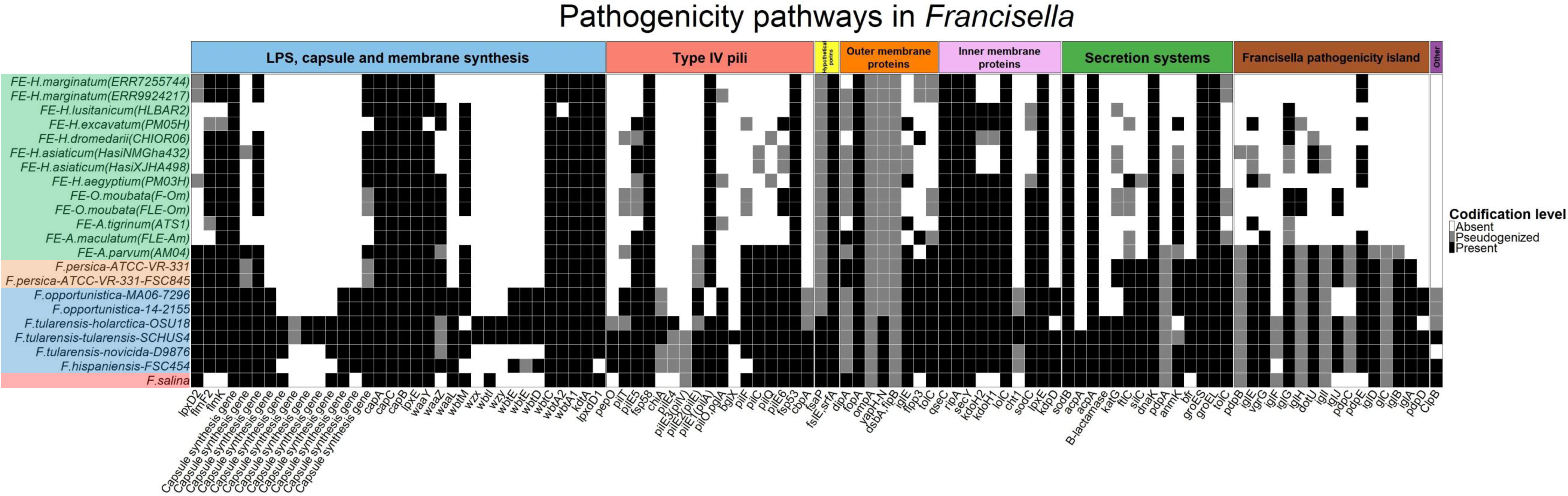
Evolution of pathogenicity-related pathways in the *Francisella* genus. The heatmap shows the conservation of genes involved in pathogenic pathways, using a color gradient where dark tones indicate intact coding sequences, lighter tones represent pseudogenes, and white indicates gene absence. Columns correspond to specific genes associated with these pathways/functions. The organisms are arranged according to the phylogenomic tree from Figure 2.

## Discussion

Ticks are key vectors of zoonotic pathogens, and their microbiome may facilitate the colonization and transmission of pathogens, including *Borrelia, Francisella, Rickettsia, Ehrlichia*, and the Crimean-Congo hemorrhagic fever virus(Baneth 2014; de la Fuente et al. 2023; Pustijanac et al. 2024). Here, we studied *Francisella* associated with ticks, reconstructing a robust phylogenomic framework that allows hypothesizing their biogeographic pattern of distribution and coevolutionary scenario with their hosts. Furthermore, we compared their predicted functional repertoire and its evolution, with a focus on vitamin biosynthesis, known to be involved in complementing host nutrition (Gerhart et al. 2016; Duron et al. 2018; Gerhart et al. 2018; Buysse et al. 2021), and virulence genes, previously characterized among *Francisella* pathogens (Farlow et al. 2005; Nano and Schmerk 2007; Degabriel et al. 2023).

### Phylogeny, Coevolution, and Biogeography

The phylogenomic relationships among FE strains and within the whole *Francisella* genus were resolved with maximum statistical support (Figure 2, left panel). The FE strains were recovered as monophyletic, consistent with previous studies (Duron et al. 2018; Gerhart et al. 2018), but supported here with denser taxon sampling. FE and FI strains from a sister group, both deriving from FP strains. Notably, pathogenic strains were recovered as paraphyletic, suggesting that the FE and FI originated from apathogenic lineage (Figure 2, left panel). This is consistent with previous studies that suggested that the endosymbiotic condition of *Francisella* evolved from a pathogenic ancestor (Gerhart et al. 2016; Duron et al. 2018; Gerhart et al. 2018). *Francisella persica*, hosted by *Argas arboreus* was indicated as a possible intermediate evolutionary stage (Suitor and Weiss 1961; Larson et al. 2016). Our phylogenies (Figure 2, left panel; Figure 3; Figure S3) are consistent with this hypothesis, placing *F. persica* (FI) as the sister clade of FE strains, with both phylogenetically derived within the pathogenic *Francisella* clade. Genomic comparisons are also partially consistent with this view, since *F. persica* shares several features with FE strains, such as genome size, coding genes, and vitamin biosynthesis, as well as some pathogenicity-related pathways shared with FP strains. At the same time, a closer inspection of the genomic diversity within the FE clade suggests a more complex scenario.

The results revealed partial co-cladogenesis between FE and their tick hosts. While host-symbiont phylogenies at the tick family level show some inconsistencies (Figure 2, dotted boxes), a higher degree of congruence emerges at the genus level (Figure 2). In the genus *Amblyomma*, phylogenies match completely, differently from a previous study using MLST data (Binetruy et al. 2020). In the case of *Hyalomma*, our result displays almost full co-cladogenesis, with only the nodes of FE-*H*. *dromedarii* (Figure 2, left) and *H. dromedarii* resulting incongruent with the respective host phylogeny (Figure 2, right). These findings point to evolutionary stable FE-tick associations, potentially shaped by short-term coevolutionary dynamics (e.g., within genus), as well as deeper symbiont transfers across unrelated ticks’ lineages (Bontemps et al. 2024). These transfers or jumps between hosts, which may be the product of ancient geographic colonization events of ticks’ lineages.

MLST genes from FE (Binetruy et al. 2020) were used to expand the sampling of FE strains across various tick species (Figure 3). The resulting phylogenetic tree aligns with the phylogenomic tree (Figure 2, left panel), showing FL and FP strains as the earliest divergences, with FI and FE as sister lineages (complete tree in Figure S3), supporting previous hypotheses (Duron et al. 2018). Historical distribution of ticks and horizontal transfer events of FE lineage could explain the observed co-cladogenesis, considering the timing and phylogenetic patterns of *Amblyomma* and *Hyalomma* genera (Sands et al. 2017; Uribe et al. 2024). On one hand, a recent study has postulated the origin of diversification of *Amblyomma* (around 37 million years ago –mya–, with height posterior density (HPD) between 40 and 26 mya), potentially fostered by the faunistic flow in the final Antarctic Bridge connection between Australia and South American lands (Uribe et al. 2024). Later, the phylogenetic pattern of *Amblyomma* indicates a posterior colonization to Indomalayan/Afrotropical/Australasia ecoregions close to the Oligocene-Miocene boundaries (node IAA, figure 1 in (Uribe et al. 2024)). At that moment, a massive biodiversity dispersion occurred through Bering Land Bridge (BLB) between Nearctic to Palearctic ecoregions (Jiang et al. 2019), driven by the expansion of the boreotropical forests to the Northern Hemisphere because of the warm and humid global climate, which served as a bridge for the warm-adapted taxa (Wolfe 1978). On the other hand, the origin of diversification of main lineages of *Hyalomma* was hypothesized in the Oligocene in Eurasia (including Indomalaya), with a successive colonization event towards Afrotropical ecoregions in the Miocene through Gomphotherium landbridge (closure of Tethys Sea) (Sands et al. 2017). This last event could have been led by the progressive and intermittent collision among Africa, Arabia, and Anatolia plates (Rögl 1999) and by the aridification of the African Palearctic and Afrotropic regions (Akhmetiev et al. 2005). Our phylogenetic reconstruction of FE fits with such biogeographic history of their host genera. On one hand, the most derived American *Amblyomma* species in the MLST tree are mostly those that reach a Nearctic distribution (Figure 3), species that could have undergone a colonization through the BLB until the Indomalaya realm. In Indomalaya region, both tick genera could have been living in sympatry, which may have supported the horizontal spread of *Francisella*, facilitated during co-feeding on the same vertebrate hosts (Duron, Sidi-Boumedine, et al. 2015) and the ability of bacteria to colonize tick salivary glands (Klyachko et al. 2007; Budachetri et al. 2014; Wright et al. 2015).

Overall, these findings highlight the geographic and phylogenetic diversification of FE strains in their tick hosts and regions, contributing to a deeper evolutionary understanding and providing the first biogeographical framework for FE. However, further research on FE in ticks, including genomes of FEs hosted by the early divergent species of *Hyalomma*, and more specimens from Indomalaya, Afrotropic and Australasia ecoregions regions, could provide deeper insights into biogeographic and coevolutionary patterns, as well as horizontal transfer paths of these symbionts.

### Genome evolution

Comparative genomics analysis of metabolic pathways between FE, FI, FP, and FL strains revealed some common traits but also notable differences. FE genomes are markedly smaller than those of FP and FL (Table S2 and Figure S1). This is consistent with the notion that genome reduction and its associated changes are well-documented phenomena in symbionts of ticks (Gerhart et al. 2016; Buysse et al. 2021; Nardi et al. 2021; Duron 2024) and other eukaryotes (McCutcheon and Moran 2010; Castelli et al. 2024). In FE, the higher presence of pseudogenes as compared to other *Francisella* is indicative of a relatively recent, and possibly ongoing genome reduction (Figure S2) (Sjödin et al. 2012; Gerhart et al. 2016; Duron et al. 2018; Gerhart et al. 2018).

Regarding FE and FI, it is notable that, despite their genomic erosion, they retain a high conservation in certain functional categories, such as J (Translation, ribosomal structure and biogenesis), C (Energy production and conversion), T (Signal transduction mechanisms), D (Cell cycle control, cell division, chromosome partitioning), and F (Nucleotide transport and metabolism) (Figure 4). This suggests that, despite experiencing genome reduction and gene erosion, these bacteria preserve key pathways necessary for survival and functionality as endosymbionts (Keeling and McCutcheon 2017; Wernegreen 2017; McCutcheon et al. 2024). Conversely, they are markedly depleted in other functions that are probably accessory/dispensable in the tick host, such as E (amino acid transport and metabolism) and S (function unknown). This pattern of genomic evolution in FE is confirmed by analyses at the level of single COGs, showing that the repertoire of FE (and FI) is largely a subset of those of FP and FL, with only few private COGs (Figure 5), thereby corroborating that extensive genomic erosion is the main cause of those differences within the genus.

Focusing specifically on key functions for members of *Francisella* genus, such as vitamin biosynthesis and pathogenicity, our results broadly confirmed expectations, adding novel relevant insights. Overall, FE retain the vast majority of vitamin biosynthetic pathways present in FL and FP (Figure 6). These include heme, which cannot be synthesized by ticks, depending on vertebrate blood as an exogenous source (Perner et al. 2016; Oleaga et al. 2017; Oleaga et al. 2021), and possibly on their endosymbionts during off-host stages (Stavru et al. 2020). This idea is supported by the localization of FE symbionts in the Malpighian tubules (Duron et al. 2018; Gerhart et al. 2018), which are involved in converting hemolymph into essential vitamin products (Sonenshine D.E & Roe 2013). Conversely, a significant exception is the set of NAD synthesis genes, which are all but one absent/pseudogenized in all FE, indicating the absence of a functional pathway, differently from FI, which display an almost equivalent repertoire to the FP (Figure 6). This suggests that ticks and their FE rely on other sources for NAD.

The lack of a portion of genes for pantothenate/CoA synthesis in FE and FI suggest that these bacteria may get the corresponding pathways intermediate metabolites via their tick hosts. Additionally, while biotin plays a key role in central metabolism (Tong 2012; Sirithanakorn and Cronan 2021), and was proposed as a nutrient provision for ticks by their symbionts (Duron and Gottlieb 2020; Buysse and Duron 2021), we did not find a full conservation of its biosynthesis among FE, given that several genes are eroded in the most derived FE hosted by *Hyalomma* spp. This could indicate another instance of “singularity” of nutritional supplementation needs among different ticks, as already suggested for members of the genus *Ixodes* (Duron 2024). Curiously, the synthesis of pyridoxal-phosphate (B6), a cofactor for numerous enzymes (Bender 1994; Leklem 2001), while stably present among FE and in FI, shows erosion in the pathogenic *F. opportunistica* strains. Taken together, these patterns confirm that ticks rely on their FE endosymbionts for B-vitamin provisioning (Braz et al. 1999; Duron 2024) and are consistent with a potential intermediary stage for FI between FE and FP (Suitor and Weiss 1961; Gerhart et al. 2018).

For what concerns virulence genes, the pattern of gene erosion observed in FE strains (Figure 7) points towards an endosymbiotic adaptation (Gerhart et al. 2016; Duron et al. 2018; Gerhart et al. 2018). As a Gram-negative bacteria, pathogenic members of the genus *Francisella* feature an outer membrane rich in lipopolysaccharides (LPS) and outer membrane proteins, and even a capsule, with these components playing key roles in virulence and host immune interactions (Lederberg 1956; Lüderitz et al. 1982; Nikaido 2003; Napier et al. 2012). While some of those core structural components are retained also in FE (and consistently in FI), several others are indeed eroded. Similarly, secretion systems, essential for transporting molecules across the membrane (Costa et al. 2015), are notably reduced in FE and in FI, yet remain intact in FP and FL strains. Some multifunctional genes, such as the core stress-response gene dnaK, vital for the Type VI secretion system (T6SS) (Alam et al. 2020) but also for thermal tolerance, are nevertheless preserved. The presence/absence patterns of the genes within the *Francisella* Pathogenicity Island (FPI), critical for virulence (Nano et al. 2004; Hager et al. 2006; Nano and Schmerk 2007) as well as of the type IV pili machinery, required for motility and DNA uptake (Tomaso et al. 2005; Chakraborty et al. 2008), show some evolutionarily noteworthy trends. Indeed, while eroded in most FE, they are present at an equivalent degree, and only slightly lower than in pathogens, in the FE of *A. parvum* (AM04) and in FI. This apparent intermediate stage was expected for FI but seems notable for FE of *A. parvum* (AM04), which in this regard is more akin to FI than to its closer FE relatives. Taking also into account the total number of coding genes (Supplementary table S1), these data hint towards an independently ongoing genome reduction among the different sub- lineages of the clade including FE and FI, with largely but not fully consistent results, due to the common selection pressures exerted by the interaction with ticks. This would explain the intermediate lineage-specific instances that are not fully consistent with phylogenetic relationships. This pattern is reminiscent of the evolution of other bacterial lineages with ancestral host association, such as *Coxiella* symbionts of ticks (Nardi et al. 2021). An alternative but non-mutually exclusive hypothesis is that FE of *A. parvum* (AM04) could have received the additional pathogenicity genes by horizontal gene transfer (possibly from another *Francisella*), considering that these were found encoded on a possibly plasmidic contig.

## Conclusions

This study reports six newly sequenced genomes of *Francisella* endosymbionts (FE) from ticks, adding, for the first time, five fully circularized genomes. This genomic dataset significantly enhances our understanding of FE evolution within the genus *Francisella* and their associations with tick hosts. Phylogenomic analyses revealed the presence of multiple distinct FE lineages, displaying partial signals of co-cladogenesis with their respective tick hosts, particularly within the genera *Amblyomma* and *Hyalomma*. Also, the evolutionary hypotheses allow us to trace for the first time a biogeographic framework of FE, suggesting further association with the ticks and their ancient potential distributions followed by a geographical spread and by fortuitous horizontal transfer between ticks.

From a metabolic and functional perspective, our findings support the hypothesis that FE contribute to the nutritional physiology of their tick hosts, particularly through the provision of essential vitamins. In contrast to their pathogenic relatives, FE consistently lack key virulence-associated genes, underscoring their functional shift from vertebrate pathogens to mutualistic symbionts.

According to our analyses, similarly to other nutritional symbionts of ticks, multiple FE lineages, as well as their close relative FI, are undergoing independent processes of genome reduction. These trajectories show convergent, though not entirely overlapping, patterns of gene loss, as particularly evidenced by the retention of select pathogenicity genes in both FI and the FE of *A. parvum* (AM04).

These findings provide novel insights into the evolutionary dynamics of symbiosis in tick-associated bacteria and offer a refined framework to study host–microbe coevolution in medically and veterinary relevant vector systems. However, further research, including broader genome sampling of FE, transcriptomic analyses, and functional experiments, will be essential to corroborate and expand upon these results.

## Acknowledgements

Many thanks to Hydra (https://researchcomputing.si.edu/high-performance-computing-cluster) and Drago (https://aic.csic.es/supercomputador-drago/) supercomputers, where the high-performance computing was conducted. This work was supported by the Comunidad de Madrid through the Atracción de Talento contract of JEU (REFF 2019-T2/AMB-13166); and the project GARES-2023, which was funded by Europe Community Union with the Plan de Recuperación, Transformación y Resiliencia, Nextgeneration EU (ASO and FV).

